# Dynamic alterations of retinal EphA5 expression in retinocollicular map plasticity

**DOI:** 10.1101/433904

**Authors:** Qi Cheng, Mark D. Graves, Sarah L. Pallas

## Abstract

The topographically ordered retinocollicular projection is an excellent system for studying the mechanism of axon guidance. Gradients of EphA receptors in the retina and ephrin-As in the superior colliculus (SC) pattern the anteroposterior axis of the retinocollicular map, but whether they are involved in map plasticity after injury is unknown. Partial damage to the caudal SC at birth creates a compressed, complete retinotopic map in the remaining SC without affecting visual response properties. Previously, we found that the gradient of ephrinA expression in compressed maps is steeper than normal, suggesting an instructive role in compression (Tadesse et al., 2013). Here we measured EphA5 mRNA and protein levels after caudal SC damage in order to test the hypothesis that changes in retinal EphA5 expression occur that are complementary to the changes in collicular ephrin-A expression. We find that the nasotemporal gradient of EphA5 receptor expression steepens in the retina and overall expression levels change dynamically, especially in temporal retina, supporting the hypothesis. This change in receptor expression occurs after the change in ephrin-A ligand expression. We propose that changes in the retinal EphA5 gradient guide recovery of the retinocollicular projection from early injury. This could occur directly through the change in EphA5 expression instructing retino-SC map compression, or through ephrinA ligand signaling instructing a change in EphA5 receptor expression that in turn signals the retinocollicular map to compress. Understanding what molecular signals direct compensation for injury is essential to developing rehabilitative strategies and maximizing the potential for recovery.

## Introduction

Understanding the mechanisms of compensatory neural plasticity after perinatal brain injury is important for promoting maximal functional recovery from traumatic brain injury (TBI). The retinotectal (*aka* retinocollicular in mammals) topographic map has been a valuable system for studying compensatory plasticity of topographic map formation, first in amphibians and fish and then in neonatal hamsters (Sperry, 1963; Gaze and Sharma, 1970; Yoon, 1971; Yoon, 1976; Schmidt, 1978; Udin and Schneider, 1981; Udin and Gaze, 1983; Constantine-Paton and Ferrari-Eastman, 1987). Retinocollicular map compression occurs after unilateral partial (PT) lesion of the superficial layer of caudal superior colliculus on the day of birth (Finlay et al., 1979; Wikler et al., 1986; Pallas and Finlay, 1989). This occurs without appreciable loss of retinal ganglion cells; an SC lesion of about 50% produces less than a 10% increase in retinal ganglion cell death rates (Wikler et al., 1986). Compensatory retino-SC map compression has been studied at the network and cellular levels, but the molecular mechanisms underlying map compression are unknown.

The complementary gradients of EphA receptors along the nasotemporal axis in retina and of ephrinA ligands along the rostro-caudal axis in optic tectum/SC provide a molecular basis for ordered retinocollicular topography across vertebrate species (Cheng and Flanagan, 1994; Cheng et al., 1995; Drescher et al., 1995; Nakamoto et al., 1996; Davenport et al., 1998; Feldheim et al., 1998; Frisen et al., 1998; Goodhill and Richards, 1999; Becker and Becker, 2000; King et al., 2003; Feldheim et al., 2004; King et al., 2004; Lemke and Reber, 2005; Triplett and Feldheim, 2012; Cang and Feldheim, 2013; Godfrey and Swindale, 2014; Willshaw et al., 2014). Positioning of retinal axon termination zones occurs through repulsive interactions between ephrinA ligands and EphA receptors, with axons responding to relative rather than absolute levels of retinal EphA signaling (Brown et al., 2000; Hansen et al., 2004; von Philipsborn et al., 2006; Fiederling et al., 2017), perhaps by comparing ligand levels between growth cone filopodia.

Regulation of ephrinA/EphA signaling has been shown to occur after optic nerve injury (Knoll et al., 2001; Rodger et al., 2001), and spinal cord injury (Willson et al., 2002; Irizarry-Ramirez et al., 2005). We found previously that ephrinA expression in SC is regulated after central target damage as well. In animals that had been subjected to neonatal damage to caudal SC, resulting in retinocollicular map compression, a steepened ephrinA5 gradient was seen (Tadesse et al., 2013). The increase in gradient steepness was accompanied by a transient reduction in expression of both ephrin-A2 and -A5. How EphA receptors might be involved in the re-establishment of topography after central target damage has not been studied. Thus in the present study we asked whether matching changes might occur in retinal EphA receptor expression after neonatal caudal SC ablation (PT), and if so, whether EphA expression may be regulated in a way that would predict map compression. We chose EphA5 because it is expressed in a gradient in the retina (Feldheim et al., 1998) and has been shown to be important in retinocollicular mapping (Feldheim et al., 2004). Retinal EphA5 mRNA and protein levels were examined in animals with partial SC ablation using quantitative real time (q) PCR and Western blots, respectively. We found that both the protein and mRNA expression levels of retinal EphA5 in PTs were increased at P5, and that there was a steeper gradient of EphA5 expression along the nasotemporal axis of the retina. The changes in EphA5 expression occurred after the damage-induced changes in ephrin-A ligand expression reported previously (Tadesse et al., 2013). Taken together, our results raise the possibility that the steepening of the ephrin-A/EphA5 gradients instructs the compensatory neural plasticity resulting in retinotopic map compression, preserving visual function after perinatal traumatic brain injury.

## Materials and Methods

### Animals

Syrian hamsters (*Mesocricetus auratus*) of different postnatal ages were used in this study. This animal model was chosen for two main reasons. One is that Syrian hamsters have a short gestation time (15.5 days) and are born at an earlier stage of retinal axon outgrowth than other rodents (Schneider, 1973). Secondly, our previous examinations of retinocollicular map plasticity were conducted in hamsters (Pallas and Finlay, 1989; Huang and Pallas, 2001; Razak et al., 2003; Tadesse et al., 2013). All animals were bred and cared for in the Georgia State University Animal Facility. They were maintained on a 14:10 light/dark cycle, and were given hamster chow and water *ad libitum*. All procedures used on the animals in this study met or exceeded standards of humane care developed by the National Institutes of Health and the Society for Neuroscience and were approved by the Institutional Animal Care and Use Committee.

### Neonatal surgery

In order to initiate retinocollicular map compression, caudal lesions of the superficial gray layer of the superior colliculus were made at birth (P0), with the aim to remove approximately half of the SC. All lesions were done under sterile conditions. Anesthesia was induced with 4% isoflurane gas/oxygen (0.5 L/min) and maintained throughout the surgery at 1-2%. Respiration rate and withdrawal reflexes were monitored continuously. The skull over the midbrain was exposed and the superficial layer of the caudal part of the right SC was ablated using a heat cautery applied briefly to the skull. The incision was closed with 6-0 silk or VetBond^®^ (3M, St. Paul, MN). The pups were then taken off the anesthetic, injected with subcutaneous fluids (Lactated Ringer’s solution + 5% dextrose at 25cc/kg), given a drop of doxapram (2mg/kg) sublingually to stimulate respiration, and rewarmed. They were closely monitored for 3 days after the surgery, and given subcutaneous fluids (Lactated Ringer’s solution + 5% dextrose at 25cc/kg) and carprofen (5-10 mg/kg). As a control for the effects of the specific location of the brain damage, some pups received a lesion of right frontal cortex rather than SC. The surgical procedures were otherwise identical.

### Quantification of gene expression

In order to obtain tissue from the retina for quantitative real-time PCR (qPCR), animals were first deeply anesthetized with sodium pentobarbital (150mg/kg, IP). After all withdrawal reflexes ceased, orbits were removed and both retinae were dissected free. Retinae were then dissected free from the pigmented epithelium and carefully divided into temporal and nasal halves (as defined by the position of the medial and lateral rectus muscles) under RNase-free conditions. Total RNA from each half of each retina was isolated with an RNeasy Isolation Kit according to manufacturer directions (Qiagen Inc., Valencia, CA). SuperScript Reverse Transcriptase II (Invitrogen Corp, Carlsbad, CA) was used for cDNA synthesis from total RNA, according to the manufacturer’s protocol. The concentrations of total RNA and cDNA were measured in triplicate with a BioPhotometer (Eppendorf Instruments Inc., Westbury, NY). An ABI PRISM 7700 Fast Real Time PCR System was used for qPCR (comparative C_T_ method, *aka* 2-(DeltaDeltaC(T)) method; Applied BioSystems, Foster City, CA) (Schmittgen and Livak, 2008). TaqMan Gene Expression Assays (Cat. No: 4351372, Applied BioSystems, Foster City, CA) provided predesigned probes and primers for our gene of interest, and TaqMan^®^ Fast Universal PCR Mix (Applied BioSystems, Foster City, CA) was used in PCR reactions. The amount of cDNA added to the PCR reactions was 50-100ng as determined by primer efficiency, and the amount of cDNA in each primer reaction was identical within each PCR reaction. Three to eight individual animals were used in each experimental group. PCR reactions were run in triplicate at 95°C for 20 sec and 60°C for 30 sec, followed by 30 cycles at 95°C for 30 sec. A negative control omitting hybridization probes was also run in triplicate to verify PCR specificity. Relative levels of EphA5 mRNA in the retina were calculated by normalizing to expression levels of the housekeeping gene GAPDH. GAPDH expression remained stable in a previous study of traumatic brain injury (Cook et al., 2009), making it an appropriate internal control for qPCR in this study. Comparison of EphA5 expression levels with GAPDH levels allowed correction for variations in total RNA and/or cDNA synthesis between samples. In order to compare RNA expression differences between the normal and lesioned group, measurements of the threshold cycle number (C_T_) were determined by analysis of the quantitative qPCR reaction using ABI Prism 7700 SDS software. C_T_ indicates the point in the PCR reaction at which the amount of amplified target gene sequence reaches a fixed threshold value related to its abundance, providing a determination of target concentration relative to a reference gene (in this case GAPDH). Differences between the normal and the PT group were analyzed using a Student’s *t*-test. All results are expressed as mean ± standard error of the mean (SEM). A probability value (p) of <0.05 or less was considered statistically significant.

### Western blot analysis

In order to obtain retinal tissue for Western blots, the animals were deeply anesthetized with sodium pentobarbital (150mg/kg weight, IP). Retinae were rapidly dissected, immediately frozen on dry ice, and then stored at −80°C. The tissue was immersed in lysis buffer containing 1% Triton X-100, 0.1% SDS, 150nM NaCl, 0.01M Sodium Phosphate (pH 7.4), 2mM EDTA, and a protease inhibitor cocktail (Roche, Indianapolis, IN USA) (Matsunaga et al., 2000), mechanically homogenized, and centrifuged for 25min in order to obtain the supernatant. The concentration of solubilized protein was determined with a Bio-Rad Protein Assay kit (Bio-Rad, Hercules, CA USA). The retinal lysate with loading buffer was boiled for 3 min prior to loading onto an 8% sodium dodecyl sulfate (SDS)-polyacrylamide gel with the same amount of protein (5ug) in each well. The proteins in the gel were transferred onto nitrocellulose membranes (Bio-Rad, Hercules, CA). Equivalent transfer of proteins in each well was confirmed in all cases by Ponceau red staining. Each membrane was incubated in a blocking solution of 5% nonfat dry milk in phosphate buffer with 0.1% Tween 20 for 1 hr at room temperature, followed by primary antibody (anti-rabbit EphA5 1:250, R&D Systems Inc., Minneapolis, MN) diluted in blocking buffer, and was then incubated overnight at 4°C. After washing 3 times with TBST buffer (TBS buffer in 1% Tween 20), the membranes were incubated in secondary antibody (biotinylated goat anti-mouse, Bio-Rad, Hercules, CA) diluted 1:2000 in blocking buffer for 30 min. Membranes were then washed in TBST. Protein signals were detected using a chemiluminescent substrate kit (Thermo Scientific, Rockford, IL USA) according to the manufacturer’s protocol. The films were scanned and optical density measurements of the bands were performed using ImageQuant TL7.0 (GE Healthcare, Creve Coeur, MO USA) using the same settings in each case.

### Retinal axon tracing and image analysis

In order to compare axonal outgrowth in SC between normal and lesioned animals, we used axon labeling to visualize the axons in SC. The animal anesthesia procedure was the same as that used in the neonatal surgery. A 0.5ul solution of cholera toxin β-Alexa Fluor 594 (CTB-594, 2mg/ml in PBS, Molecular Probe) was injected through the eyelid and into the posterior chamber of the left eye using a glass pipette tip connected to a Hamilton syringe (Hamilton Co., Reno, NV). Hamster pups were euthanized at different ages (P1, P3, and P5) by deeply anesthetizing with sodium pentobarbital (150mg/kg, IP), and were then intracardially perfused with PBS containing 4% paraformaldehyde (PFA). The head was immersed overnight in 4% PFA before the SC was dissected out for whole mounts. The images of CTB labeled retinal axons in the whole mount of each SC were captured with an AxioCam Hrm digital camera (Carl Zeiss Optical, Thornwood, NY) through a Nikon (Melville, New York) fluorescence microscope with a 4x objective. This allowed us to visualize the dye front from above in the SC as a whole. This labeled area was measured with AxioZeiss Version 3.0 software (Carl Zeiss Optical, Thornwood, NY). The background fluorescence of the unlabeled (left) SC was useful in determining the boundaries of the SC in the labeled (right) SC and in distinguishing background fluorescence from that in the labeled axons. Both rostrocaudal distance grown and proportion of the SC occupied by the fluorescing axons were calculated for normal and lesioned cases. Three to six individual animals were used in each experimental group. All results are expressed as mean ± SEM. The statistical differences between the normal and the PT animals were analyzed using a Student’s *t*-test. A probability value (p) of 0.05 or less was considered a significant difference.

## Results

In a previous study, we demonstrated that damage to the caudal part of SC at birth affected expression of ephrin-A2 and –A5 in the SC in concert with retino-SC map compression (Tadesse et al., 2013). The ligand expression gradient became steeper in proportion to the degree of compression of the map, raising the possibility that the change in ligand gradient instructs compression of the retinotopic map in SC. Surprisingly, the amount of ephrinA was significantly and specifically downregulated in the lesioned SC. In this study, we addressed three questions raised by the previous study in regards to the effect of partial target loss on expression of the EphA5 receptor in the retina of animals with compressed retinocollicular maps. One was whether the slope of the EphA5 receptor gradient would be altered to match the altered slope of the ligand gradient. The second question was whether there would be receptor compensation for the change in ligand expression levels. Third, if retinal EphA receptors are affected by SC target loss, we wanted to establish the timing of the changes; that is, whether it is the ligand or the receptor expression that changes first after SC injury.

Syrian hamsters were used in this study (Finlay et al., 1979; Pallas and Finlay, 1989). In total, 122 animals were used, including 53 normal, 6 sham control and 63 PT (neonatal partial ablation of caudal SC) animals (Table 1). Most data was collected at postnatal ages (P)1 to P5, with the first 24 hours after birth designated as P0. This time period corresponds to the point when expression levels of ephrinAs are highest and when the retinal axons are invading the SC. In addition to the data from P1-P5, 6 animals at P8 and 6 at P15 (3 Normal, 3 PT each) were added to examine the decline in EphA5 with age.

**Table 1.**

| **Postnatal**<br>**Age**<br>**(P)** | **Number of Normal**<br>**Animals** |  |  | **Number of**<br>**Cortical**<br>**Lesion**<br>**Animals** | **Number of**<br>**SC Lesion Animals** |  |  |
| --- | --- | --- | --- | --- | --- | --- | --- |
|  | Axonal<br>Labeling | qPCR | Western<br>Blot | qPCR | Axonal<br>Labeling | qPCR | Western<br>Blot |
| P1 | 3 | 4 | 3 | - | 4 | 4 | 3 |
| P3 | 5 | 9 | 7 | - | 6 | 9 | 4 |
| P5 | 4 | 8 | 4 | 6 | 5 | 13 | 9 |
| P8 | - |  | 3 |  | - |  | 3 |
| P15 | - |  | 3 |  | - |  | 3 |
| Total | 12 | 21 | 20 | 6 | 15 | 26 | 22 |

### Retinal EphA5 expression during early postnatal development

Retinal EphA5 expression has been well studied during embryonic development. EphA5 is expressed in embryonic retinae in mice (Feldheim et al., 1998; Cooper et al., 2009) and chick (Cheng et al., 1995). In order to assess the normal pattern of retinal EphA5 expression during early postnatal development in the hamsters, we examined the time course of changes in retinal EphA5 expression during early visual development using qPCR. We found that expression levels of retinal EphA5 mRNA changed after birth (P1, 0.8 ± 0.03 log_10_(EphA5/GAPDH), n=3), to a peak at P3 (1.2 ± 0.06, n=3; 50% increase from P1 value, p < 0.005, *t*-test), followed by a decrease at P5 (1.0 ± 0.10, n=3; decline to 25% of P3 value, p < 0.0001, *t*-test), then declining to very low levels by P8 and P15 (P8: 0.038 ± 0.02, n=3; P5 vs. P8 p < 0.001, *t*-test. P15: 0.044 ± 0.008, n=3; P5 vs P15 p < 0.001) (**Fig. 1**). This time course is similar to that seen previously with ephrinA ligand expression (Tadesse et al., 2013). Determining that EphA5 is expressed in hamster retina during early postnatal development allowed us to assess how retinal EphA5 expression is regulated with age in animals that had early damage to the SC.

**Figure 1.**
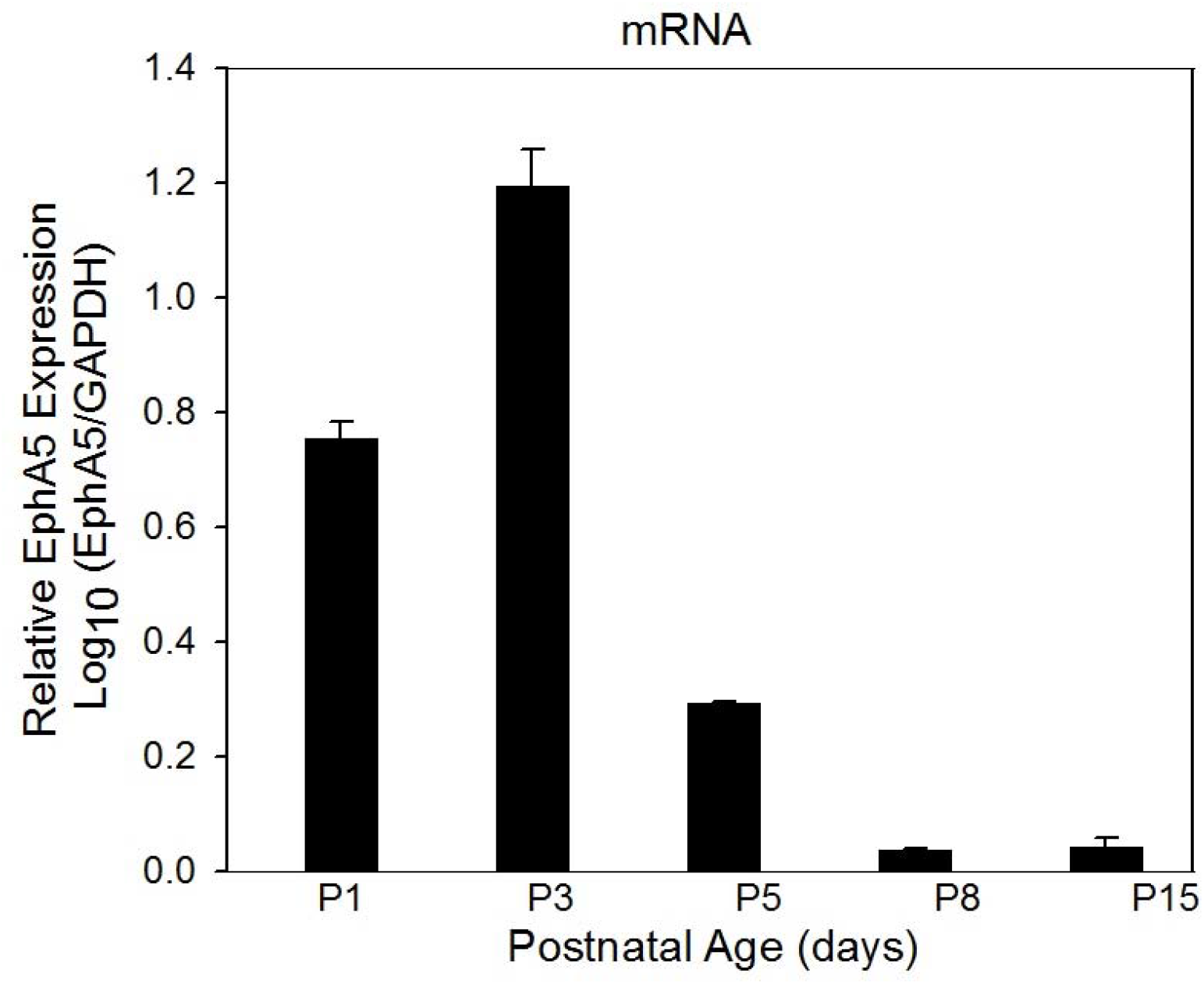
Postnatal development of retinal EphA5 expression pattern. Retinal EphA5 mRNA levels were quantified with qPCR. Relative expression levels of retinal EphA5 mRNA increased after birth, achieved a peak at P3, and then decreased by P5. By P8-15, EphA5 mRNA expression was markedly decreased.

### Outgrowth of RGC axons in SC is influenced by SC damage at birth

In order to address whether damage to central targets of retinal ganglion cells (RGCs) would influence the rate and pattern of axon outgrowth in SC, we compared the projection pattern of retinocollicular axons in SC of lesioned animals with that in SC of normal animals by using intraocular injection of CTB for anterograde tracing of the retino-SC projection. Alexa-fluor 594-conjugated cholera toxin B-subunit (CTB) was injected into the vitreous chamber of the left eye on the day of birth (P0), and whole-mounts or coronal sections of the SC were prepared for analysis of labeled retinal axons at P1, P3, and P5. (Note that P1 is the day after birth.) In the normal animals, the retinal axons had already extended across the rostral half of SC by P1, had reached the far caudal SC by P3, and had covered the entire SC by P5. In contrast, in animals with partial (PT) lesions in caudal SC, the retinal axons had invaded only the rostral third of the remaining SC by P1 and only the rostral two-thirds by P3. By P5, however, the retinal axons covered most of SC. In order to compare the developmental progression of retino-SC innervation in the two groups in a quantitative fashion, we measured the area of SC that was innervated by the retinal axons compared to the total SC area. We found that the proportion of SC innervated by retinal axons in PT animals was significantly lower than that in normal animals on all 3 days (PT cases at P1:16.6 ± 4.01%, n=4; P3: 73.5 ± 2.82%, n=6; P5: 85.4 ± 4.10%, n=5; Normal cases at P1: 72.9 ± 0.98%, n=3; P3: 94.9 ± 1.51%, n=5; P5: 99.6 ± 0.56%, n=4; Student’s *t*-test, p<0.05, **Fig. 2A**). We also measured the distance that the axons grew from the rostral-most edge along the rostrocaudal axis of SC to provide an indication of growth rate. We found that the distance grown in PT animals (P1: 0.3 ± 0.04mm, n=4; P3: 1.1 ± 0.05mm, n=6, P5: 1.3 ± 0.09mm, n=5) was significantly less from birth to P5 compared with the distance grown in SC of normal animals (P1: 1.2 ± 0.05mm, n=3; P3: 1.7 ± 0.04mm, n=5; P5: 1.8 ± 0.03mm, n=4; Student’s *t*-test, p<0.005, **Fig. 2B**). The slopes of the lines in Figure 2B suggest that retinal axons in the lesioned animals were initially growing more rapidly than in the normal animals from P1 to P3, and at about the same rate as normal from P3-5. This initial rapid outgrowth may occur to compensate for the damage to caudal SC at P0 causing a brief arrest of retinal axon outgrowth into SC. By P5 the axons have innervated nearly the entire extent of the lesioned SC. Importantly, these data show that the retinal axons in the lesioned cases are not likely to be directly affected by the P0 lesion, because their growth cones are far from the caudal SC at that point. Our previous electrophysiological studies show that the entire extent of the lesioned SC is eventually innervated (Pallas and Finlay, 1989; Huang and Pallas, 2001). Taken together, these data suggest that retinal axon growth rate can be dynamically influenced by damage to central targets, raising the possibility that the factors responsible for organizing axon outgrowth in the retina may be changed to compensate for damage to the SC target.

**Figure 2.**
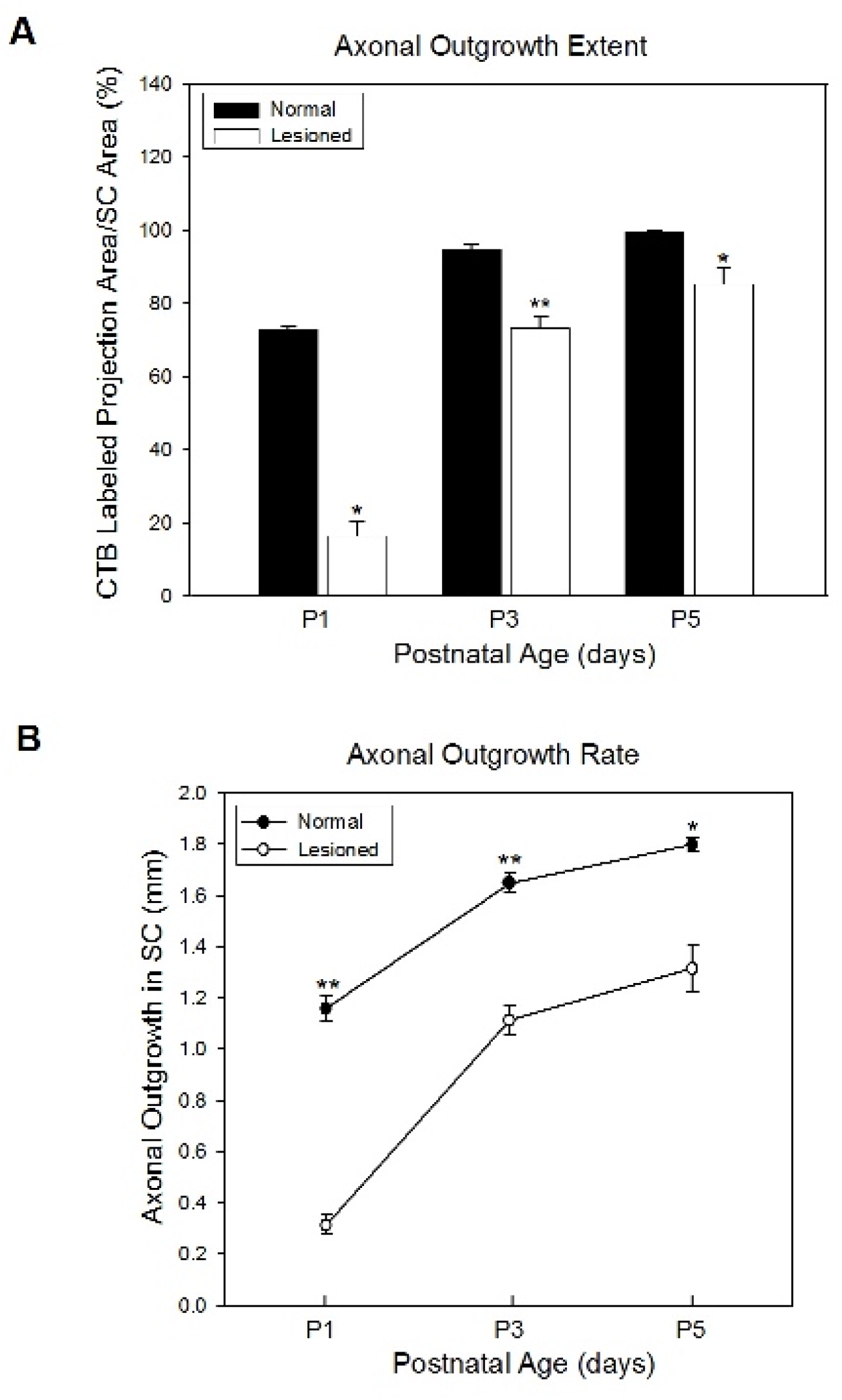
Outgrowth of retinal axons into SC is affected by partial lesion of SC at birth. Following CTB injection into the eye at birth, images of CTB labeled axons in SC in the normal and lesioned animals were analyzed at different postnatal ages in normal and PT lesioned animals. (A) The proportion of SC occupied by retinal axons in PT animals was significantly smaller than that in normal animals at all 3 ages, but was especially low at P1, 24hr after the damage. (B) The actual distance grown by retinal axons along the rostro-caudal axis of SC was greater in normal than in PT lesioned animals because the lesioned SC is smaller. The axonal outgrowth rate between P1 and P3 was greater in PT than in normal animals; although the growth rate between P3 and P5 in PTs was similar to that in normal animals. *indicates p<0.005, ** indicates p<0.001 (Student’s *t*-test). Error bars show mean ± SEM. Each group contains 3-6 animals.

### Retinal EphA5 expression is altered after SC damage at birth

In our previous study, we demonstrated that the expression of ephrin-A2 and –A5 in the SC was changed after caudal SC lesion at birth (Tadesse et al., 2013). The ligand expression gradient became steeper in proportion to the degree of compression of the map, raising the possibility that the change in ligand gradient instructs compression of the retinotopic map in SC. We also found that the amount of ephrin was significantly and specifically downregulated in the lesioned SC. The drop in ephrin-A expression in SC occurred first in caudal SC, followed by a drop in expression in rostral SC.

In order to measure whether expression of retinal EphA5 receptors is also altered after the partial target loss, we compared EphA5 mRNA and protein expression levels in the retina of normal animals to expression in animals that had undergone partial SC ablation at birth. We used qPCR to assess EphA5 mRNA levels and Western blots to measure EphA5 protein levels in the retinae at P5 when ephrin-A ligand levels are highest. Only the left retina was used, which in the PT lesioned group is the retina contralateral to the lesion and thus most directly affected, given the small amount of binocular overlap in this species. β-actin was used as a comparison to control for non-specific effects of the lesions on gene and protein expression. We confirmed that the retinal β-actin mRNA expression (**Fig. 3A**), and retinal β-actin protein levels (**Fig. 3B and C**) did not differ significantly across conditions. The relative qPCR analysis revealed that the EphA5 mRNA levels in the retina of PT animals were significantly increased (ratio of target gene to GAPDH gene abundance, 0.7 ± 0.15 log_10_ (EphA5/GAPDH), n=5) compared to normal (0.1 ± 0.06, n=5) (Student’s *t*-test, p<0.05, **Fig. 3A**). A 5-fold increase in EphA5 protein in PT animals (5073.3 ± 95.00 optical density (OD) units, n=3) compared to normal conditions (1099.7 ± 542.70 OD units, n=3) was observed with Western blot analysis (Student’s *t*-test, p<0.005, **Fig. 3B and C**). These data suggest that a specific upregulation of EphA5 can be induced by the SC lesion. The upregulation may be a compensatory response to the lesion and/or to the previously described reduction in ligand levels.

**Figure 3.**
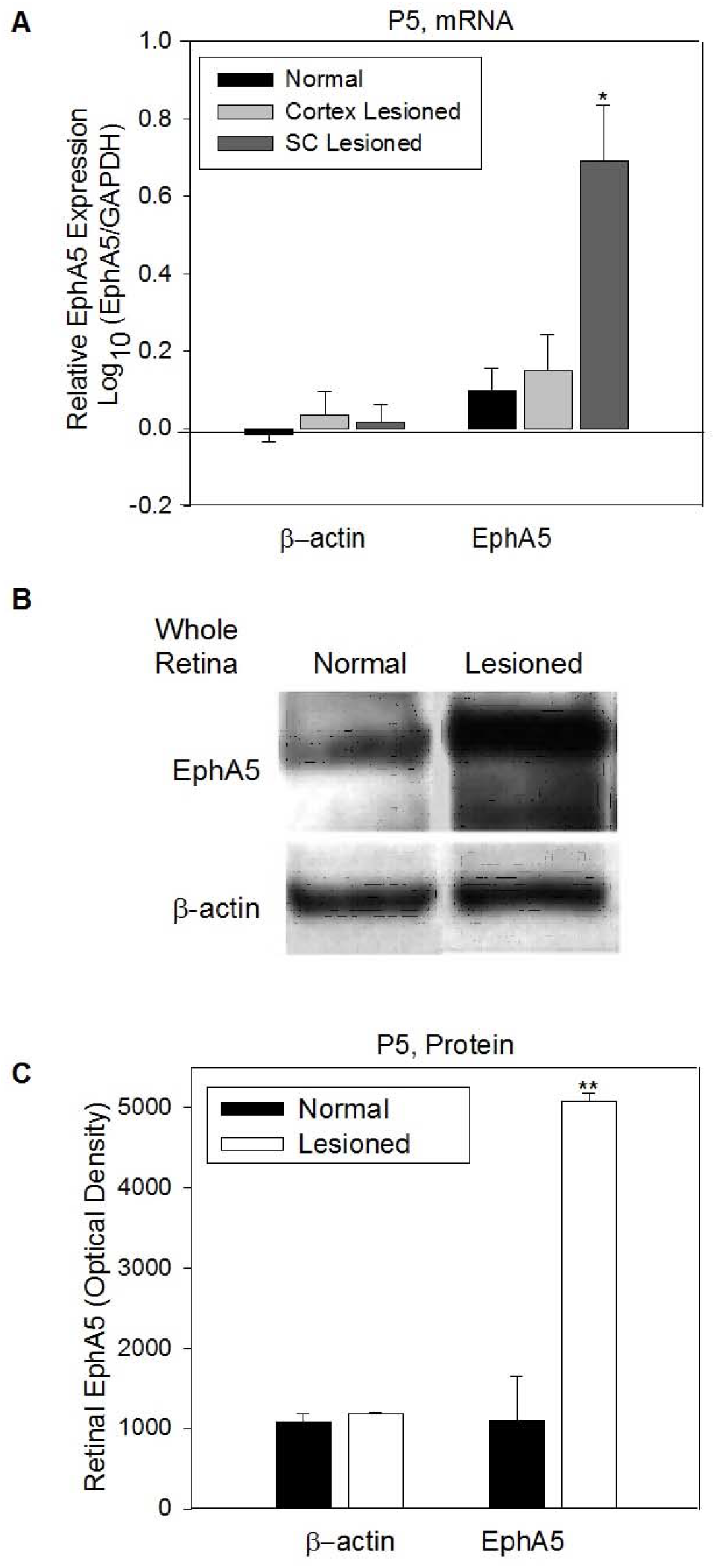
Changes in retinal EphA5 expression in SC-lesioned animals are specific. (A) Quantification of EphA5 and β-actin gene expression in intact retina by qPCR. Retinal EphA5 mRNA levels were up regulated by P5 in PT animals but not in animals with cortical lesions or in normal animals. (B) Retinal EphA5 and β-actin protein levels were assessed by Western blot analysis of the whole retina. EphA5 was increased in PT animals although β-actin was unchanged. (C) Western blot quantification. EphA5 protein was increased 5-fold in PT animals compared with that in normal animals but β-actin protein levels were unaffected by the lesion. *indicates p<0.005. Error bars show mean ± SEM.

In order to control for the possibility that these apparently compensatory changes in retinal EphA5 expression after partial SC lesion might instead be a non-specific response to brain damage in general, we compared EphA5 and β-actin mRNA expression levels in P5 hamster retinae obtained from normal and PT lesioned animals to that in animals with lesions of frontal cortex. The gene expression levels of both β-actin and EphA5 in cortex-lesioned animals (0.2 ± 0.09, n=6) did not differ significantly from that in normal animals (0.1 ± 0.06, n=5) (Student’s *t*-test, p>0.05, **Fig. 3A**). The data suggest that the changes in retinal EphA5 expression seen in PT cases are specific to that pathway and not a general effect of any surgical procedure or of lesions in another region on EphA5 expression.

To address whether the changes in EphA5 expression seen in the above analysis of the entire retina as a whole affected the nasotemporally-increasing EphA5 gradient, we divided each hamster retina into nasal and temporal halves using the positions of the medial and lateral rectus muscles on the eye as landmarks. EphA5 expression was then measured in each of the retinal halves and compared between the normal and lesioned animals at two postnatal ages corresponding to the time points taken in our ephrinA study (Tadesse et al., 2013). We found that at P3, EphA5 mRNA expression in temporal retina of PT animals (−0.3 ± 0.27 log_10_(EphA5/GAPDH), n=4) was significantly decreased compared to that in normal animals (0.4 ± 0.09, n=4; Student’s *t*-test, p<0.05, **Fig. 4A**). Furthermore, at P3 in PT animals, there was a 12 fold decline in EphA5 protein levels in the temporal retina, and a 5 fold decline in nasal retina compared with normal animals (PT: OD relative to background in temporal retina 230.7±98.23; OD relative to background in nasal retina 336.2 ± 115.25, n=3; Normal: temporal: OD 2801.4 ± 99.68; nasal: OD 1674.3 ± 38.43; Student’s *t*-test, p<0.001) (**Fig. 4B, C**). Two days later, at P5, the EphA5 mRNA levels in PT lesioned animals (temporal: 0.6 ± 0.07; nasal: 0.4 ± 0.05, n=8) were increased by approximately four fold above normal in nasal retina, and by three fold in temporal retina compared with normal animals (temporal: 0.20 ± 0.0002; nasal: 0.09 ± 0.0001, n=3; Student’s *t*-test, p<0.05) (**Fig. 4D**). Protein expression showed changes comparable to those seen with mRNA expression; we found that EphA5 protein levels in retinae of lesioned animals (temporal: OD 2249.0 ± 232.04; nasal: OD 998.300 ± 125.63, n=3) were increased by 2.5 fold above normal in nasal retinae (OD 454.19 ± 87.86, n=3) and by 4 fold above normal in temporal retinae (OD 565.400 ± 231.67, n=3; Student’s *t*-test, p<0.05) (**Fig. 4E, F**). The data show that the initial decrease in retinal EphA5 expression in lesioned compared to normal animals at P3 has been reversed by P5 in both nasal and temporal retina. These data support the hypothesis that target damage can delay maturation of EphA5 receptor expression in the retinal axons, and show that there is a time-dependent, dynamic effect of the lesions on expression levels that can affect the receptor gradient as it does the ligand gradient (Tadesse et al., 2013).

**Figure 4.**
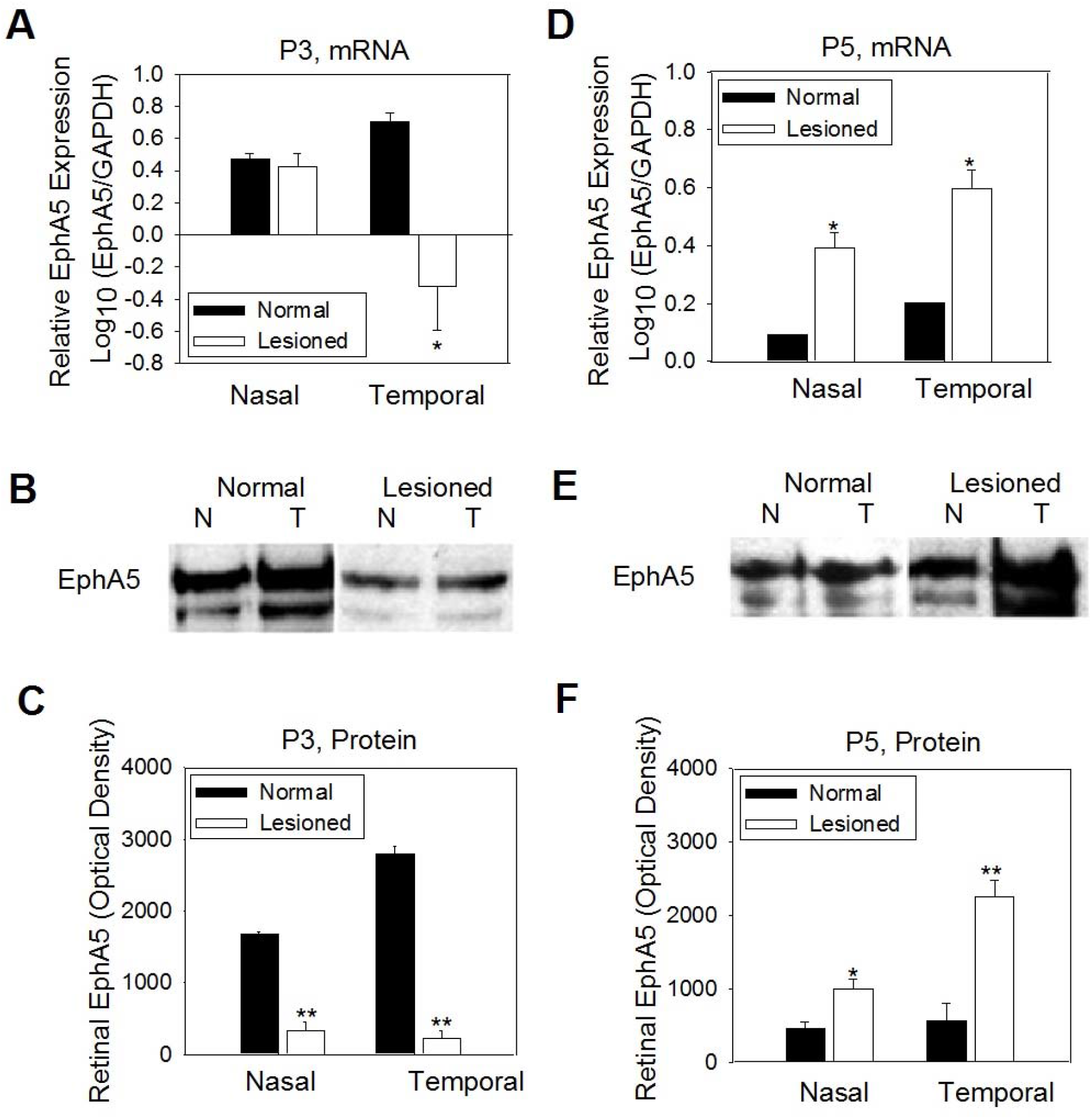
Retinal EphA5 receptor expression changed in a time-dependent, dynamic fashion after partial lesion of SC at birth. Retinal EphA5 mRNA or protein expression levels in nasal and temporal retina were examined and quantified by qPCR or Western blot, respectively. (A) At P3, EphA5 mRNA expression was significantly decreased below normal in temporal retina of PT animals. (B) A representative Western blot of retinal EphA5 protein expression in a normal and a PT case at P3. N: Nasal, T: Temporal. (C) Western blot quantification. At P3, both temporal and nasal retinal EphA5 protein levels were significantly decreased in the PT cases compared with those in normal animals. (D) In contrast, at P5, EphA5 mRNA levels of both nasal or temporal retina were significantly increased above normal in PT animals. (E) Both temporal and nasal retinal EphA5 proteins at P5 were increased in lesioned cases compared with normal cases. N: Nasal, T: Temporal. (F) Western blot quantification. EphA5 protein in nasal and temporal retina was significantly increased at P5 in PT cases. * indicates p<0.05, ** indicates p<0.001 (Student’s *t*-test). All error bars are indicate mean ± SEM.

### The retinal EphA5 gradient is steeper during retino-SC map compression

Our main interest was to determine whether retinal EphA5 might be involved in the retinal axon behavior that results in compensatory retino-SC map compression after target damage. One way in which this could occur is if the EphA gradient was compressed in reaction to the caudal SC lesion. In our previous study we found that the ephrinA gradient in caudally-lesioned SC is compressed, suggesting that the EphA5 gradient might also be steeper than normal after the caudal SC lesion (Tadesse et al., 2013). To test this hypothesis, we compared EphA5 mRNA and protein expression gradients in nasal and temporal retinae in normal and PT animals. We defined gradient steepness as the ratio between the EphA5 mRNA expression levels in temporal versus nasal retina. Consistent with the dynamic changes in expression levels seen from P3 to P5, we found that the steepness of the EphA5 gradient changed during this time period. The EphA5 mRNA gradient was transiently reversed at P3 in PT animals because the reduction in expression at this time point occurs in the temporal retina first and is more extreme than in the nasal retina (cf. **Fig. 4A**). This change resulted in an approximately 5-fold decrease in the temporal/nasal ratio (−3.7±1.57, n=3) compared to normal animals (1.2±0.30, n=3) (Student’s *t*-test, p<0.05, **Fig. 5A**). Similarly, the steepness of the EphA5 protein expression gradient was reduced by 13-fold (−1.1 ± 3.42, n=3) such that the gradient was nearly eliminated at P3 in the lesioned animals compared to normal animals (11.3±0.66, n=3) (Student’s *t*-test, p<0.001, **Fig. 5B**). In marked contrast to the P3 findings, at P5 the gradient of EphA5 mRNA expression in PT animals (4.1 ± 1.39, n=8) was not only restored but was steeper by approximately 2-fold compared to normal (2.2 ± 0.007, n=3) (**Fig. 5C**). Similarly, the steepness of the EphA5 protein expression gradient at P5 was increased by 6 fold (normal: 0.5 ± 0.23, PT: 6.4 ± 0.23, n=3) (Student’s *t*-test, p<0.001, **Fig. 5D**). The changes in the steepness of the EphA receptor gradient in PT animals suggest that these guidance molecules may be involved in the retinal axon behavior that compensates for the target damage through SC map compression.

**Figure 5.**
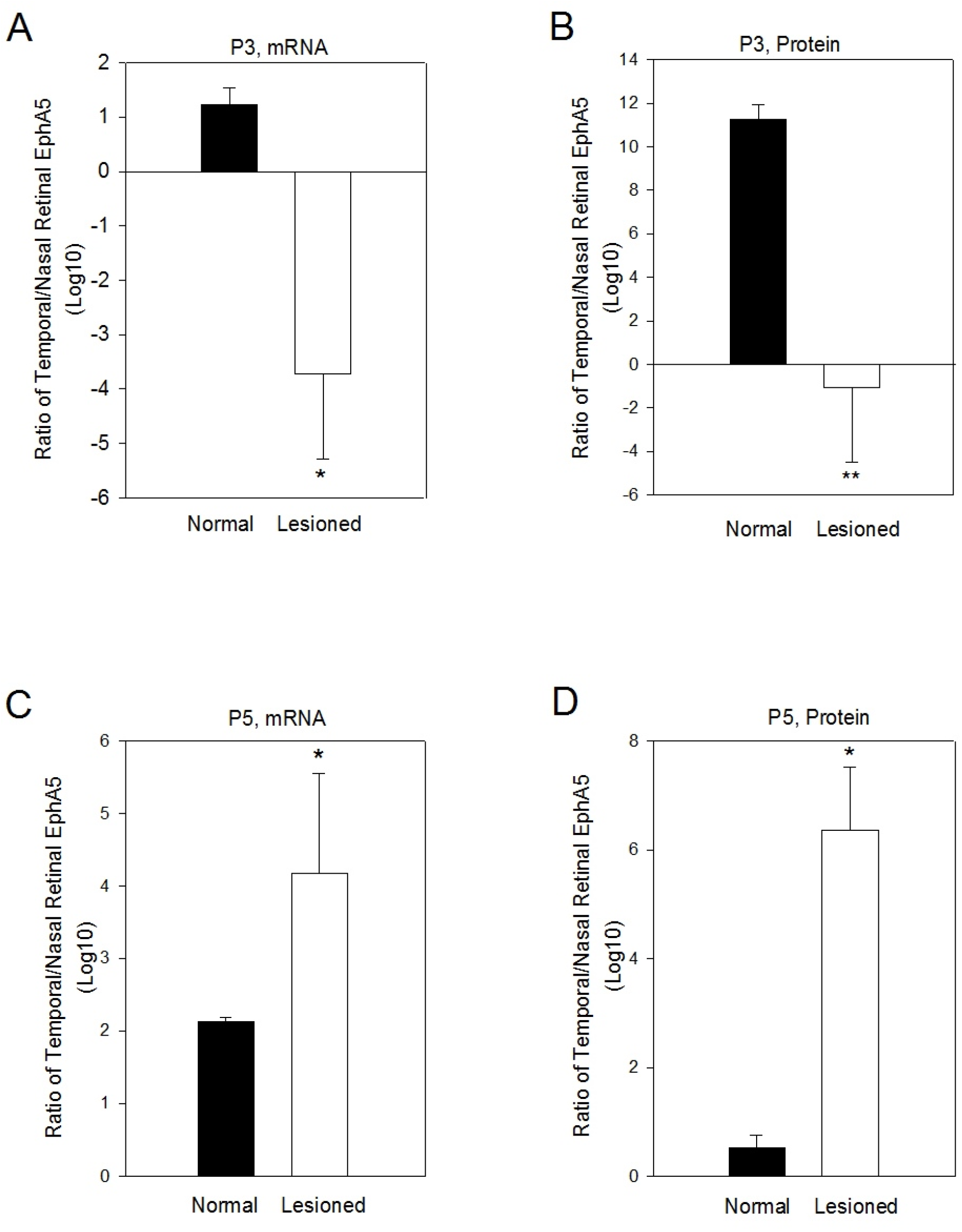
The retinal EphA5 gradient is steeper after partial lesion of SC at birth. EphA5 gradient steepness, defined as the temporal/nasal ratio of retinal EphA5 expression levels in retina, was increased by P5 in lesioned animals. (A) The EphA5 mRNA expression gradient was reversed at P3 in PT cases, with nasal retina expressing at a higher level than temporal retina. (B) The EphA5 protein expression gradient was also reversed, by 13 fold. (C) At P5, the temporal/nasal ratio of EphA5 mRNA expression was higher in PT animals by approximately 2 fold compared to normal. (D) The temporal/nasal ratio of EphA5 protein level was significantly higher in PT cases than in normal animals (*p<0.05; **<0.001, Student’s *t*-test). All error bars represent mean ± SEM.

### Lesion-induced changes in retinal EphA5 expression occur after changes in ephrinA expression in SC

In a previous study, we demonstrated that the amount of ephrinA was significantly and specifically downregulated by P1 in the lesioned SC, within 24 hr after partial SC lesion at birth (Tadesse et al., 2013). If the changes in ephrinA expression in the lesioned SC cause the changes in retinal EphA5 expression, we would predict that they would occur first. To test this prediction, we examined the time course of changes in receptor expression within 15 days after the lesion of SC using qPCR. Retinal EphA5 expression in lesioned animals at P1 (0.8 ± 0.15 log_10_ (EphA5/GAPDH), n=3) was not significantly different from that in normal animals (0.8±0.03, n=3). By P3 in lesioned animals, EphA5 mRNA was transiently downregulated (0.2 ± 0.29, n=3) compared with that in normal animals (1.2 ± 0.06, n=3) (Student’s *t*-test, p<0.05). By P5, EphA5 mRNA (1.0 ± 0.10, n=3) was significantly upregulated compared with normal P5 animals (0.3 ± 0.003, n=3) (Student’s *t*-test, p<0.005). By the second week of postnatal development, retinal EphA5 mRNA expression in lesioned animals was much reduced (P8: 0.08 ± 0.02, n=3; P15: 0.026 ± 0.01, n=3) and was not significantly different from that in normal animals (P8: 0.038 ± 0.02, n=3; P15: 0.044 ± 0.008; p> 0.05; normal data are taken from Fig. 1) (**Fig. 6**). Thus, the changes in EphA5 expression occur prior to major developmental events necessary to formation of a complete retino-SC map (Clancy et al., 2001; 2007). Taken together with the previous observation (Tadesse et al., 2013) that ephrinA3 and A5 expression in SC was decreased by P1, 24 hr after the partial lesion of SC, these results indicate that the retinal EphA5 expression is altered after the ephrinA expression changes in lesioned animals. These data raise the possibility that the lesion-induced changes in collicular ephrinA5 expression instruct the changes in retinal EphA5. Alternatively, the target damage may trigger changes in both ligand and receptor, with the receptor changes taking a longer time to be manifested. In either case, our findings in this study strongly suggest an involvement of the EphA/ephrinA system in compensation for target damage leading to retino-SC map compression.

**Figure 6.**
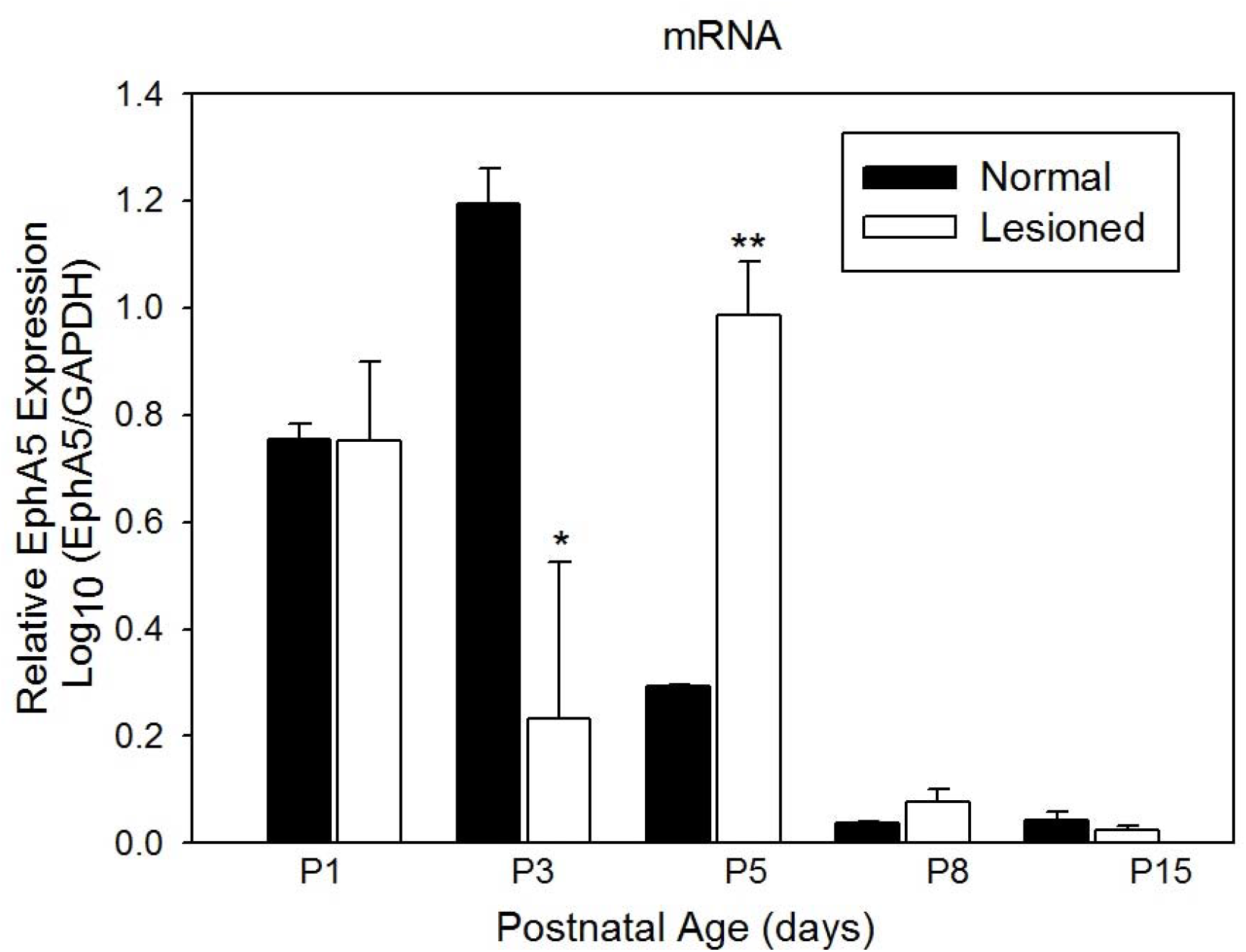
Lesion-induced changes in retinal EphA5 expression. Retinal EphA5 mRNA was quantified with qPCR from animals between 1 and 15 days after birth. At P1, EphA5 mRNA expression in PT animals was not significantly different from that in normal animals. Expression was significantly decreased from normal at P3, and then was increased above normal by P5 in PT animals, declining markedly and returning to normal levels in the second postnatal week. * indicates p<0.05 (Student’s *t*-test). Error bars are presented as mean ± SEM.

## Discussion

Traumatic brain injury is one of the major causes of death and disability in young people, making it a serious public health problem (Yue et al., 2013). Thus, understanding the mechanisms of recovery from perinatal brain damage is essential to developing treatment strategies (Zink, 2001). The progress of basic and clinical research on retinal repair and regeneration has been hampered by an inability to achieve efficient and accurate target innervation in adults (Crair and Mason, 2016; Lim et al., 2016; Bray et al., 2017). Understanding how neonatal retinal ganglion cells can accomplish this task is likely to lead to innovations in treatment for adults.

The ephrinA/EphA family of repulsive axon guidance cues is required for development of topographic maps in the visual pathway (Feldheim et al., 2004; Cang et al., 2005; see Clandinin and Feldheim, 2009, for review). Previous evidence from our lab demonstrated that a change in ephrinA gradient steepness occurs after caudal SC lesion at P0 (Tadesse et al., 2013). This suggested a causal role in map compression, and predicted a complementary change in EphA receptor expression levels. Therefore, the hypothesis tested here was that a correlation exists between the map compression resulting from caudal SC damage at birth, changes in ephrinA levels, and changes in EphA levels. Our data demonstrate that the retinal EphA5 expression gradient for mRNA and protein levels changes dynamically during the recovery phase and becomes steeper than normal during early postnatal development in PT animals. They show that the steepening of the retinal EphA5 gradient is correlated with the compensatory neural plasticity that results in compression of the retinocollicular map after injury and partial target loss. Together with our previous findings on changes in ephrinA ligand expression in SC after partial SC injury at birth (Tadesse et al., 2013), our data suggest that lesion-induced changes in retinal EphAs could play an instructive role in recovery from neonatal brain injury.

### Methodological considerations

An important consideration for interpreting our results concerns the effect of the P0 lesions on ingrowing retinal axons (Fischer et al., 2017). Retinal axons in neonatal rats can grow across lesion sites (Symonds et al., 2001) but likely behave differently than spared axons. We argue based on the results herein that the lesions are unlikely to damage retinal axons directly because axon outgrowth is limited to very rostral SC at P0 when lesions are made (see also Jhaveri et al., 1991; Chen et al., 1995; Ding et al., 2001). A more detailed, single axon analysis of retinal axon responses to SC damage is underway.

We measured mRNA and protein levels in the retina itself, and assume that these are reflective of protein levels at the terminal arbor, where ligand binding occurs. How a signal from the damaged region of SC may be communicated to the retinal axon terminals and then to the nuclei in the retina to alter gene expression is not known and will be addressed in future experiments. Our results caution that ephrinA gene knockout in mice may alter EphA expression in a compensatory fashion. This could have unpredictable effects due to promiscuity of the receptors for several ligands.

### Central target damage alters retinal EphA5 receptor expression levels in a time-dependent fashion

Development of EphA5 expression has been studied during embryonic development in chicks and mice (Cheng et al., 1995; Feldheim et al., 1998) and postnatally in mice (Cooper et al., 2009). We provide in this study the first demonstration of the postnatal expression pattern of EphA5 mRNA and protein in Syrian hamsters. Hamsters have been the subject of many studies of developmental plasticity in the visual system (Schneider, 1973; Rhoades and Chalupa, 1978; Finlay et al., 1979; Emerson et al., 1982; Pallas and Finlay, 1989), in large part because they are born at an earlier stage of brain development than other common lab rodents (Clancy et al., 2001; Nagarajan et al., 2010).

In order to assess whether EphA5 expression is affected by partial SC lesion at birth, we used qPCR to compare expression in normal and lesioned animals. We found that retinal EphA5 expression was downregulated by the third day after lesion of SC, and was then upregulated by P5, 6 days after the lesions. The downregulation of EphA5 gene expression could be an initial response to the injury of the SC target. The decline of EphA5 levels at P3 could indicate either that the lesion of caudal SC triggers an expression-inhibiting signal, or that it interrupts an expression-promoting factor. The possible outcome of the decline is not clear but it may result in a temporary loss of sensitivity of retinal axons to ephrin ligand in the target, such that they would not be influenced by the ephrin gradient during the initial recovery phase. The ensuing upregulation of EphA5 expression by P5 would complement the reduction in ephrin ligand levels in SC that are induced by damage (Tadesse et al., 2013). The increase in EphA5 levels by P5 after the lesion may be important for SC map compression, and may compensate for ligand expression changes in SC, although whether there is a cause and effect relationship is not known. In sum, the dynamic changes in EphA5 expression following injury to the SC target suggest that the endogenous EphA receptors play an important role in the re-establishment of retinocollicular topography, and may be required for compression of the SC map.

Studies of optic nerve regeneration in adults have also reported changes in EphA expression. For example, 1 month after optic nerve section in adult rats, a decrease in EphA5 expression was found (Rodger et al., 2001). In contrast, an increase in EphA5 expression was correlated with the restoration of topography after optic nerve injury in goldfish (King et al., 2003). These results suggest that the effect of visual pathway damage on EphA expression is highly dependent on where the damage occurs and on time after occurrence. Our results showing that alteration of retinal EphA5 expression levels occurs in a time-dependent fashion stress the importance of examining changes in expression over time after injury. The finding that EphA5 expression is regulated by methylation (Petkova et al., 2010) may provide insight into the mechanism underlying injury-induced changes. Our results argue that EphA5 plays an important role in functional recovery from injury in the visual pathway. Whether damage-induced alterations in expression of retinal EphA5 receptors and their ephrinA5 ligands in SC cause SC map compression, and if so, how, is currently under study (Balmer et al., 2017).

### The steeper EphA5 gradient is correlated with SC map compression

The retinal EphA gradient is required for ordered retinotectal/retinocollicular topography during normal visual development (Cheng et al., 1995; Brown et al., 2000; Feldheim et al., 2004; Flanagan, 2006). Our main interest in the present study was to test whether the gradient of EphA5 is altered by the SC lesion that results in retino-SC map compression. A steeper than normal gradient of EphA5 was observed in lesioned animals during early postnatal development. Our results support the hypothesis that the retinal EphA5 expression gradient changes to complement the changes in the ephrinA expression gradient after partial SC ablation at birth. Thus, EphA5 could be considered as a possible candidate for directing the re-establishment of retinocollicular topography during recovery from brain injury. Retinal axons expressing the EphA receptor can be guided by their ephrinA ligands to a particular location within the target (Walter et al., 1987; 1987), and the relative concentrations of EphA/ephrinA are related to axon growth (Brown et al., 2000; Hansen et al., 2004). Studies using genetic knock-in of EphA have demonstrated that relative, not absolute levels of EphA in neighboring RGCs determine the appropriate termination site in SC and that axons can adapt to differing levels of these guidance cues (Brown et al., 2000; Hansen et al., 2004; Reber et al., 2004; Fiederling et al., 2017). We propose that increases in relative levels of EphA5 in individual axons may facilitate axons in finding their appropriate position in the remaining collicular space by a competitive mechanism, compressing the topographic map on the remaining SC area in reaction to the steeper gradient of retinal EphA expression. This is similar to the process that occurs in establishment of normal topography (Frisen et al., 1998; Feldheim et al., 2000; Feldheim et al., 2004).

The receptive field properties of SC neurons, such as size tuning, velocity tuning, and receptive field size, are preserved in compressed SC maps (Finlay et al., 1979; Pallas and Finlay, 1989). This preservation occurs through regulation of retino-SC convergence (Huang and Pallas, 2001). Thus, understanding the compensatory mechanisms resulting in SC map compression may be instructive for understanding the mechanism of functional recovery from brain injury (Du et al., 2007; Bolsover et al., 2008).

### Relative timing of changes in EphA expression in SC map compression

The relationship between changes in retinal EphA receptor expression and changes in expression of their ligands in SC was another focus of this study. Interaction between EphA receptors and their ligands is necessary for axon guidance during normal postnatal development (Ciossek et al., 1998). If the Eph/ephrin system is responsible for SC map compression, it could occur either through ephrinAs instructing the change in EphA5 expression or through ephrinA ligands instructing a change in EphA5 expression that in turn signals the retinocollicular map to compress. Our finding that changes in ephrin-A2 and –A5 expression occur prior to changes in retinal EphA5 expression in lesioned animals (Tadesse et al., 2013) suggest that the lesion-induced changes in collicular ephrinA5 expression instruct the changes in retinal EphA5. Alternatively the target damage may trigger changes in both ligand and receptor, with the receptor changes taking a longer time to be manifested. In either case, it seems likely that complementary changes in ephrinA/EphA expression gradients are involved in SC map compression. However, whether EphA5 and its ligands interact to cause SC map compression cannot be determined from these data. Loss of function studies currently underway will be useful for determining whether there is a causal link between changes in ephrinA/EphA expression and reconfiguration of the retinotopic map after early injury.

## Conclusions

In the present study, we have demonstrated that regulation of retinal EphA5 expression is induced by partial lesion of the SC at birth. The steeper gradient of retinal EphA5 that results from this regulation is correlated with both lesion-induced changes in ephrinA expression and with SC map compression, suggesting a causal role. These results may be informative in designing clinical approaches to improvement of recovery from brain injury.

## Acknowledgements

We thank Dr. Deborah Baro for the help with qPCR, and Mike Leukam for technical assistance with the SDS-PAGE gels. We also thank the staff of the GSU animal care facility for expert animal care, and members of the Pallas lab for their comments on the manuscript. This study was supported by NIH EY/MH12696, NSF IBN-0451018, the STC Program of the National Science Foundation under Agreement No. IBN-9876754, and the Brains and Behavior Program at GSU.

## References

Balmer TS, Mudd DB, Mao YT, Cheng Q, Willshaw DJ, Pallas SL. 2017. Critical role for ephrinA in retinocollicular map compression demonstrated by gene knockout. In: 2017 Society for Neuroscience Annual Meeting. Washington, DC: Society for Neuroscience

Becker CG, Becker T. 2000. Gradients of Ephrin-A2 and Ephrin-A5b mRNA during retinotopic regeneration of the optic projection in adult zebrafish. J Comp Neurol 427:469.

Bolsover S, Fabes J, Anderson PN. 2008. Axonal guidance molecules and the failure of axonal regeneration in the adult mammalian spinal cord. Restorative neurology and neuroscience 26:117–130.

Bray ER, Noga M, Thakor K, Wang Y, Lemmon VP, Park KK, Tsoulfas P. 2017. 3D visualization of individual regenerating retinal ganglion cell axons reveals surprisingly complex growth paths. eNeuro 4.

Brown A, Yates PA, Burrola P, Ortuno D, Vaidya A, Jessell TM, Pfaff SL, O’Leary DD, Lemke G. 2000. Topographic mapping from the retina to the midbrain is controlled by relative but not absolute levels of EphA receptor signaling. Cell 102:77–88.

Cang J, Feldheim DA. 2013. Developmental mechanisms of topographic map formation and alignment. Annu Rev Neurosci 36:51–77.

Cang J, Kaneko M, Yamada J, Woods G, Stryker MP, Feldheim DA. 2005. Ephrin-As Guide the Formation of Functional Maps in the Visual Cortex. Neuron 48:577–589.

Chen DF, Jhaveri S, Schneider GE. 1995. Intrinsic changes in developing retinal neurons result in regenerative failure of their axons. Proc Natl Acad Sci U S A 92:7287–7291.

Cheng H-J, Nakamoto M, Bergemann AD, Flanagan JG. 1995. Complementary gradients in expression and binding of ELF-1 and Mek4 in development of the topographic retinotectal projection map. Cell 82:371–381.

Cheng HJ, Flanagan JG. 1994. Identification and cloning of ELF-1, a developmentally expressed ligand for the Mek4 and Sek receptor tyrosine kinases. Cell 79:157–168.

Ciossek T, Monschau B, Kremoser C, Loschinger J, Lang S, Muller BK, Bonhoeffer F, Drescher U. 1998. Eph receptor-ligand interactions are necessary for guidance of retinal ganglion cell axons in vitro. Eur J Neurosci 10:1574–1580.

Clancy B, Darlington RB, Finlay BL. 2001. Translating developmental time across species. Neuroscience 105:7–17.

Clancy B, Kersh B, Hyde J, Darlington RB, Anand KJ, Finlay BL. 2007. Web-based method for translating neurodevelopment from laboratory species to humans. Neuroinformatics 5:79–94.

Clandinin TR, Feldheim DA. 2009. Making a visual map: mechanisms and molecules. Curr Opin Neurobiol 19:174–180.

Constantine-Paton M, Ferrari-Eastman P. 1987. Pre- and postsynaptic correlates of interocular competition and segregation in the frog. J Comp Neurol 255:178–195.

Cook NL, Vink R, Donkin JJ, van den Heuvel C. 2009. Validation of reference genes for normalization of real-time quantitative RT-PCR data in traumatic brain injury. J Neurosci Res 87:34–41.

Cooper MA, Crockett DP, Nowakowski RS, Gale NW, Zhou R. 2009. Distribution of EphA5 receptor protein in the developing and adult mouse nervous system. J Comp Neurol 514:310–328.

Crair MC, Mason CA. 2016. Reconnecting Eye to Brain. J Neurosci 36:10707–10722.

Davenport RW, Thies E, Zhou R, Nelson PG. 1998. Cellular localization of ephrin-A2, ephrin A5, and other functional guidance cues underlies retinotopic development across species. J Neurosci 18:975–986.

Ding Y, Yao B, Lai Q, McAllister JP. 2001. Impaired motor learning and diffuse axonal damage in motor and visual systems of the rat following traumatic brain injury. Neurol Res 23:193–202.

Drescher U, Kremoser C, Handwerker C, Löschinger J, Noda M, Bonhoeffer F. 1995. In vitro guidance of retinal ganglion cell axons by RAGS, a 25 kDa tectal protein related to ligands for Eph receptor tyrosine kinases. Cell 82:359–370.

Du J, Fu C, Sretavan DW. 2007. Eph/ephrin signaling as a potential therapeutic target after central nervous system injury. Curr Pharm Des 13:2507–2518.

Emerson VF, Chalupa LM, Thompson ID, Talbot RJ. 1982. Behavioural, physiological, and anatomical consequences of monocular deprivation in the golden hamster (Mesocricetus auratus). Exp Brain Res 45:168–178.

Feldheim DA, Kim YI, Bergemann AD, Frisén J, Barbacid M, Flanagan JG. 2000. Genetic analysis of ephrin-A2 and ephrin-A5 shows their requirement in multiple aspects of retinocollicular mapping. Neuron 25:563–574.

Feldheim DA, Nakamoto M, Osterfield M, Gale NW, DeChiara TM, Rohatgi R, Yancopoulos GD, Flanagan JG. 2004. Loss-of-function analysis of EphA receptors in retinotectal mapping. J Neurosci 24:2542–2550.

Feldheim DA, Vanderhaeghen P, Hansen MJ, Frisen J, Lu Q, Barbacid M, Flanagan JG. 1998. Topographic guidance labels in a sensory projection to the forebrain. Neuron 21:13031313.

Fiederling F, Weschenfelder M, Fritz M, von Philipsborn A, Bastmeyer M, Weth F. 2017. Ephrin-A/EphA specific co-adaptation as a novel mechanism in topographic axon guidance. Elife 6:e25533.

Finlay BL, Schneps SE, Schneider GE. 1979. Orderly compression of the retinotectal projection following partial tectal ablations in the newborn hamster. Nature (London) 280:153–154.

Fischer D, Harvey AR, Pernet V, Lemmon VP, Park KK. 2017. Optic nerve regeneration in mammals: Regenerated or spared axons? Exp Neurol 296:83–88.

Flanagan JG. 2006. Neural map specification by gradients. Curr Opin Neurobiol 16:59–66.

Frisen J, Yates PA, McLaughlin T, Friedman GC, O’Leary DD, Barbacid M. 1998. Ephrin-A5 (AL-1/RAGS) is essential for proper retinal axon guidance and topographic mapping in the mammalian visual system. Neuron 20:235–243.

Gaze RM, Sharma SC. 1970. Axial differences in the reinnervation of the goldfish optic tectum by regenerating optic nerve fibers. Exp Brain Res 10:171–181.

Godfrey KB, Swindale NV. 2014. Modeling development in retinal afferents: retinotopy, segregation, and ephrinA/EphA mutants. PloS one 9:e104670.

Goodhill GJ, Richards LJ. 1999. Retinotectal maps: molecules, models and misplaced data. Trends Neurosci 22:529–534.

Hansen MJ, Dallal GE, Flanagan JG. 2004. Retinal axon response to ephrin-as shows a graded, concentration-dependent transition from growth promotion to inhibition. Neuron 42:717–730.

Huang L, Pallas SL. 2001. NMDA antagonists in the superior colliculus prevent developmental plasticity but not visual transmission or map compression. J Neurophysiol 86:1179–1194.

Irizarry-Ramirez M, Willson CA, Cruz-Orengo L, Figueroa J, Velazquez I, Jones H, Foster RD, Whittemore SR, Miranda JD. 2005. Upregulation of EphA3 receptor after spinal cord injury. J Neurotrauma 22:929–935.

Jhaveri S, Edwards MA, Schneider GE. 1991. Initial stages of retinofugal axon development in the hamster: evidence for two distinct modes of growth. Exp Brain Res 87:371–382.

King C, Lacey R, Rodger J, Bartlett C, Dunlop S, Beazley L. 2004. Characterisation of tectal ephrin-A2 expression during optic nerve regeneration in goldfish: implications for restoration of topography. Exp Neurol 187:380–387.

King CE, Wallace A, Rodger J, Bartlett C, Beazley LD, Dunlop SA. 2003. Transient upregulation of retinal EphA3 and EphA5, but not ephrin-A2, coincides with reestablishment of a topographic map during optic nerve regeneration in goldfish. Exp Neurol 183:593–599.

Knoll B, Isenmann S, Kilic E, Walkenhorst J, Engel S, Wehinger J, Bahr M, Drescher U. 2001. Graded expression patterns of ephrin-As in the superior colliculus after lesion of the adult mouse optic nerve. Mech Dev 106:119–127.

Lemke G, Reber M. 2005. Retinotectal mapping: new insights from molecular genetics. Annu Rev Cell Dev Biol 21:551–580.

Lim JH, Stafford BK, Nguyen PL, Lien BV, Wang C, Zukor K, He Z, Huberman AD. 2016. Neural activity promotes long-distance, target-specific regeneration of adult retinal axons. Nat Neurosci 19:1073–1084.

Matsunaga T, Greene MI, Davis JG. 2000. Distinct expression patterns of eph receptors and ephrins relate to the structural organization of the adult rat peripheral vestibular system. Eur J Neurosci 12:1599–1616.

Nagarajan R, Darlington RB, Finlay BL, Clancy B. 2010. ttime: an R package for translating the timing of brain development across mammalian species. Neuroinformatics 8:201–205.

Nakamoto M, Cheng HJ, Friedman GC, McLaughlin T, Hansen MJ, Yoon CH, O’Leary DD, Flanagan JG. 1996. Topographically specific effects of ELF-1 on retinal axon guidance in vitro and retinal axon mapping in vivo. Cell 86:755–766.

Pallas SL, Finlay BL. 1989. Conservation of receptive field properties of superior colliculus cells after developmental rearrangements of retinal input. Vis Neurosci 2:121–135.

Petkova TD, Seigel GM, Otteson DC. 2010. A role for DNA methylation in regulation of EphA5 receptor expression in the mouse retina. Vision Res.

Razak KA, Huang L, Pallas SL. 2003. NMDA receptor blockade in the superior colliculus increases receptive field size without altering velocity and size tuning. Journal of Neurophysiology 90:110–119.

Reber M, Burrola P, Lemke G. 2004. A relative signalling model for the formation of a topographic neural map. Nature 431:847–853.

Rhoades RW, Chalupa LM. 1978. Functional and anatomical consequences of neonatal visual cortical damage in superior colliculus of the golden hamster. J Neurophysiol 41:1466–1494.

Rodger J, Lindsey KA, Leaver SG, King CE, Dunlop SA, Beazley LD. 2001. Expression of ephrin-A2 in the superior colliculus and EphA5 in the retina following optic nerve section in adult rat. Eur J Neurosci 14:1929–1936.

Schmidt JT. 1978. Retinal fibers alter tectal positional markers during the expansion of the retinal projection in goldfish. J Comp Neurol 177:279–295.

Schmittgen TD, Livak K. 2008. Analyzing real-time PCR data by the comparative C(T) method. Nat Protoc 3:1101–1108.

Schneider GE. 1973. Early lesions of the superior colliculus: Factors affecting the formation of abnormal retinal projections. Brain Behav Evol 8:73–109.

Sperry RW. 1963. Chemoaffinity in the orderly growth of nerve fiber patterns and connections. Proc Natl Acad Sci USA 50:703–710.

Symonds ACE, Rodger J, Tan MML, Dunlop SA, Beazley LD, Harvey AR. 2001. Reinnervation of the superior colliculus delays down-regulation of ephrin A2 in neonatal rat. Exp Neurol 170:364–370.

Tadesse T, Cheng Q, Xu M, Baro DJ, Young LJ, Pallas SL. 2013. Regulation of ephrin-A expression in compressed retinocollicular maps. Dev Neurobiol 73:274–296.

Triplett JW, Feldheim DA. 2012. Eph and ephrin signaling in the formation of topographic maps. Semin Cell Dev Biol 23:7–15.

Udin SB, Gaze RM. 1983. Expansion and retinotopic order in the goldfish retinotectal map after large retinal lesions. Exp Brain Res 50:347–352.

Udin SB, Schneider GE. 1981. Compressed retinotectal projection in hamsters: fewer ganglion cells project to tectum after neonatal tectal lesions. Exp Brain Res 43:261–269.

von Philipsborn AC, Lang S, Loeschinger J, Bernard A, David C, Lehnert D, Bonhoeffer F, Bastmeyer M. 2006. Growth cone navigation in substrate-bound ephrin gradients. Development 133:2487–2495.

Walter J, Henke-Fahle S, Bonhoeffer F. 1987. Avoidance of posterior tectal membranes by temporal retinal axons. Development 101:909–913.

Walter J, Kern-Veits B, Huf J, Stolze B, Bonhoeffer F. 1987. Recognition of position-specific properties of tectal cell membranes by retinal axons in vitro. Development 101:685–696.

Wikler KC, Kirn J, Windrem MS, Finlay BL. 1986. Control of cell number in the developing visual system. II. Effects of partial tectal ablation. Dev Brain Res 28:11–21.

Willshaw DJ, Sterratt DC, Teriakidis A. 2014. Analysis of local and global topographic order in mouse retinocollicular maps. J Neurosci 34:1791–1805.

Willson CA, Irizarry-Ramirez M, Gaskins HE, Cruz-Orengo L, Figueroa JD, Whittemore SR, Miranda JD. 2002. Upregulation of EphA receptor expression in the injured adult rat spinal cord. Cell Transplant 11:229–239.

Yoon M. 1971. Reorganization of retinotectal projection following surgical operations on the tectum in goldfish. Exp Neurol 33:395–411.

Yoon MG. 1976. Progress of topographic regulation of the visual projection in the halved optic tectum of adult goldfish. J Physiol 257:621–643.

Yue JK, Vassar MJ, Lingsma H, Cooper SR, Yuh EL, Mukherjee P, Puccio AM, Gordon W, Okonkwo DO, Valadka A, Schnyer DM, Maas A, Manley GTMDPD, Casey SS, Cheong M, Dams-O’Connor K, Hricik AJ, Knight EE, Kulubya ES, Menon D, Morabito DJ, Pacheco JL, Sinha TK. 2013. Transforming Research and Clinical Knowledge in Traumatic Brain Injury (TRACK-TBI) Pilot: Multicenter Implementation of the Common Data Elements for Traumatic Brain Injury. J Neurotrauma.

Zink BJ. 2001. Traumatic brain injury outcome: concepts for emergency care. Ann Emerg Med 37:318–332.

